# Myogenic regulatory factors MyoD and Myf5 are required for dorsal aorta formation and angiogenic sprouting

**DOI:** 10.1101/2022.01.20.477095

**Authors:** Eric Paulissen, Benjamin L. Martin

**Affiliations:** Department of Biochemistry and Cell Biology, Stony Brook University, Stony Brook, NY 11794-5215

**Keywords:** MyoD, Myf5, myogenesis, cardiovascular, artery, vein, zebrafish

## Abstract

The vertebrate embryonic midline vasculature forms in close proximity to the developing skeletal muscle, which originates in the somites. Angioblasts migrate from bilateral positions along the ventral edge of the somites until they meet at the midline, where they sort and differentiate into the dorsal aorta and the cardinal vein. This migration occurs at that the same time that myoblasts in the somites are beginning to differentiate into skeletal muscle, a process which requires the activity of the basic helix loop helix (bHLH) transcription factors Myod and Myf5. Here we examined vasculature formation in *myod* and *myf5* mutant zebrafish. In the absence of skeletal myogenesis, angioblasts migrate normally to the midline but form only the cardinal vein and not the dorsal aorta. The phenotype is due to the failure to activate vascular endothelial growth factor ligand *vegfaa* expression in the somites, which in turn is required in the adjacent angioblasts for dorsal aorta specification. Myod and Myf5 cooperate with Hedgehog signaling to activate and later maintain *vegfaa* expression in the medial somites, which is required for angiogenic sprouting from the dorsal aorta. Our work reveals that the early embryonic skeletal musculature in teleosts evolved to organize the midline vasculature during development.

**Summary statement:** The myogenic transcription factors MyoD and Myf5 have a novel function in inducing the artery through regulation of Vegf.

## INTRODUCTION

During vertebrate embryonic development, the body forms progressively from the head to the tail. The posterior extension of the body is accompanied by the continuous generation and sequential segmentation of the paraxial mesoderm into structures called somites (Martin, 2021, 2016). Skeletal myogenesis occurs within the somites in a process that is dependent upon bHLH transcription factors called Myogenic Regulatory Factors (MRFs), which initiate differentiation by activating the expression of critical muscle-related genes (Coutelle et al., 2001; Hammond et al., 2007; Rudnicki et al., 1993; Weinberg et al., 1996). Two MRFs called *myod* and *myf5* are each dispensable on their own for the formation of muscle, but loss of function of both results in the complete absence of differentiated skeletal muscle (Braun et al., 1992; Coutelle et al., 2001; Hammond et al., 2007; Hinits et al., 2011; Maves et al., 2007; Rudnicki et al., 1993, 1992). MyoD and Myf5 function by dimerizing with ubiquitously expressed bHLH E-Protein partners to induce downstream myogenic targets (Ling et al., 2014). While the role of *myod* and *myf5* are well understood in the context of paraxial mesoderm specification and muscle differentiation (Coutelle et al., 2001; Hammond et al., 2007; Row et al., 2018; Rudnicki et al., 1993; Weinberg et al., 1996), it is not clear what other roles they may play during development. In adult skeletal muscle of other model organisms, differentiating myofibers secrete factors that affect surrounding tissues. Myoblasts and myofibers can secrete Vascular Endothelial Growth Factor (VEGF) to recruit blood vessels in ischemic conditions (Bryan *et al*., 2008; Renault *et al*., 2013). However, whether or not the myogenesis and Vegf relationship is significant, or even present, in the developing embryo is not known.

In zebrafish, the formation and maturation of the somites is tightly coordinated with the establishment of midline vasculature. Angioblasts originate at the ventral-lateral edge of the somites and migrate medially along the ventral edge of the somite until they reach the midline, where they differentiate into the dorsal aorta and cardinal vein through a process known a selective cell sorting (Childs et al., 2002; Fouquet et al., 1997; Herbert et al., 2009; Jin et al., 2005; Lawson and Weinstein, 2002; Williams et al., 2010; Zhong, 2005). Retinoic acid signaling induced morphogenesis of the somites plays a critical non-autonomous role in facilitating the midline angioblast migration (Paulissen et al., 2021). Additionally, somite derived *vegfaa* ligand expression activates VEGF receptors and signaling in the angioblasts to pattern the midline vasculature, with angioblasts receiving Vegfaa signal becoming the dorsal aorta, and the remaining angioblasts sorting into the cardinal vein (Casie Chetty et al., 2017; Covassin et al., 2006; Herbert et al., 2009; Jin et al., 2017; Kim et al., 2013; Lawson et al., 2002). In absence of *vegfaa* function the dorsal aorta is absent (Rossi et al., 2016). Thus, somites play a primary role in the formation of the midline vasculature through the control of angioblast migration and fate specification.

Midline derived (notochord and floor plate) Hedgehog signaling acts upon the adjacent somites to induce *vegfaa* expression (Coultas et al., 2010; Lawson et al., 2002). However, although Hedgehog receptors and Hedgehog response genes, *ptc1* and *ptc2*, are present in other tissues adjacent to the Hedgehog ligand source, such as the spinal cord, these tissues do not express Vegf ligand (Concordet et al., 1996; Koudijs et al., 2008). This indicates that *vegfaa* induction requires additional somite-specific factors. In this study, we find that *myod* and *myf5* are required for *vegfaa* induction and the formation of the dorsal aorta. Loss of *vegfaa* in *myod*/*myf5* double mutants occurs despite intact Hedgehog signaling, and results in the formation of a single midline blood vessel that expresses venous markers. Restoring VEGF signaling in *myod*/*myf5* double mutants rescues the formation of the dorsal aorta. We propose that MyoD and Myf5 work in cooperation with Hedgehog signaling to induce Vegf ligand expression in the medial somite.

## RESULTS AND DISCUSSION

### Cardiovascular defects related to loss of myogenic transcription factors MyoD and Myf5

In this study, we investigated the role of myogenic transcription factors, MyoD and Myf5, during arterial and venous formation (Hinits et al., 2009; Maves et al., 2007; Row et al., 2018). In zebrafish and murine model systems, individual loss of *myod* or *myf5* have little effect on skeletal muscle formation on their own, but combinatorial loss results in near complete absence of myogenesis (Hinits et al., 2011; Maves et al., 2007; Rudnicki et al., 1993, 1992). Given the early viability of *myod* and *myf5* double mutants in teleosts, we investigated their role during cardiovascular development.

To identify structural defects in cardiovascular development, we performed in-situ hybridization experiments of 25 hpf zebrafish embryos using pan-endothelial markers, as well as molecular markers specific to arterial or venous fates. The pan-endothelial marker, *fli1a*, showed seemingly normal vasculature. However, a more detailed view of *myod/myf5 -/-* showed an apparent loss of one of the midline blood vessels (Figure 1A and 1B, respectively; black arrows indicating the trunk vasculature). We next looked at molecular markers of the dorsal aorta and common cardinal vein, the two predominant vascular structures in the midline of the zebrafish trunk. We used the arterial specific *cldn5b* and *aqp8a* molecular markers, and observed a complete loss of arterial expression in the trunk of *myod,myf5* double mutants (Figure 1D,1F) (Casie Chetty et al., 2017). On the other hand, venous markers *dab2* and *stab1l* show normal expression in the common cardinal vein (Figure 1H,1J).

**Figure 1.**
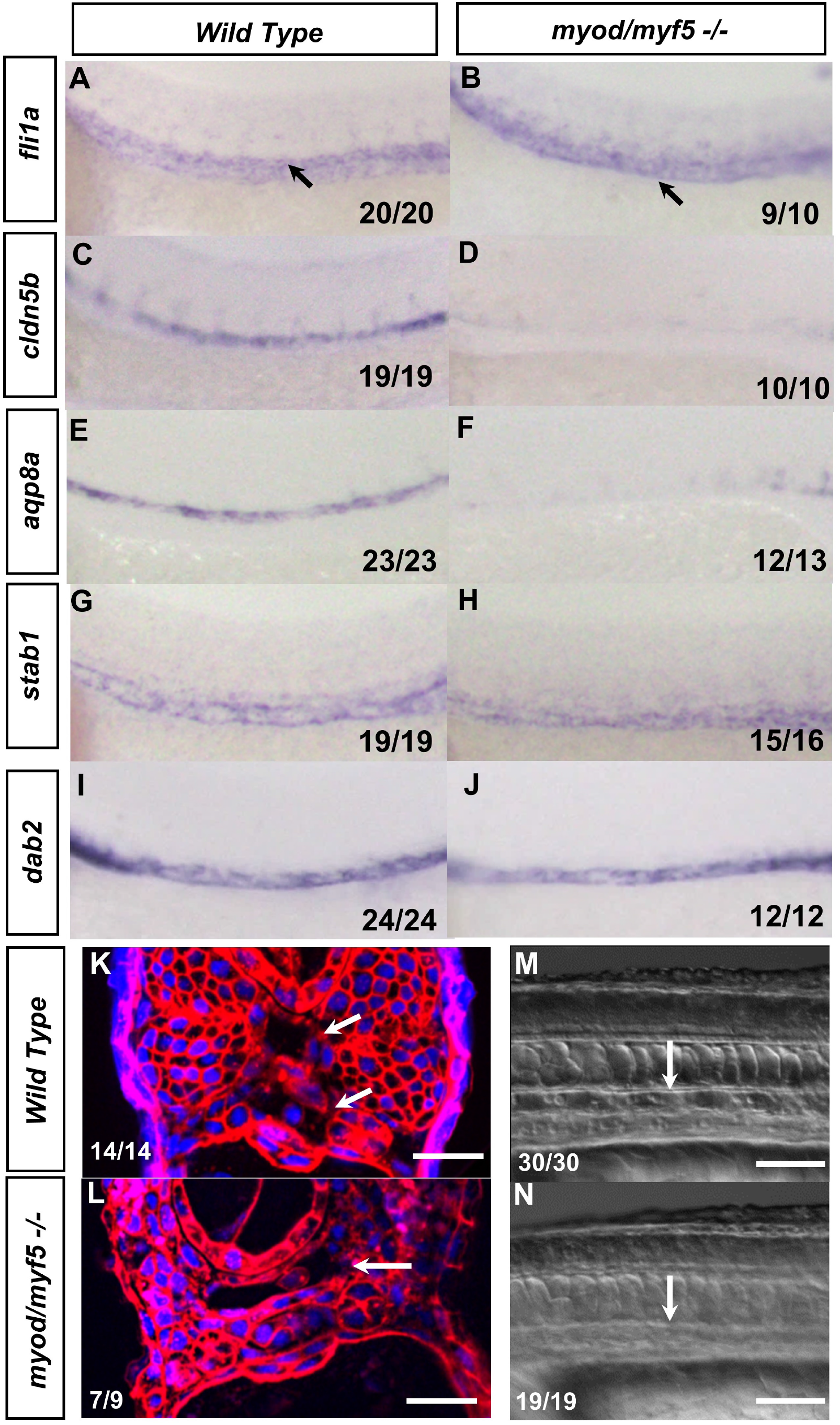
MyoD and Myf5 are required for midline arterial but not venous fate. Whole-mount in-situ hybridization of 25 hpf wild-type (**A**,**C**,**E**,**G**,**I**) and *myod/myf5 -/-* embryos (**B**,**D**,**F**,**H**,**J**). The expression of *fli1a* (pan endothelial) (**A**,**B**) *cln5b* (arterial) (**C**,**D**) *aqp8a* (arterial) (**E**,**F**) *stab1* (veinous) (**G**,**H**) and *dab2* veinous (**I**,**J**) are shown. Wild-type embryos show clear expression of both arterial markers, *cldn5b a*nd *aqp8a*, and venous markers, *stab1* and *dab2*. In *myod/myf5* double mutants, the arterial markers are largely lost (**D**,**F**), but the venous markers are normal (**H**,**J**). Fluorescent images of zebrafish trunk sections show two vessels corresponding to the artery and vein in wild-type embryos (**K**, white arrows), whereas there is a single vessel in *myod/myf5* mutants (**L**, white arrow). Analysis of circulation at 25 hpf shows normal flow in wild-type embryos and a complete absence in *myod/myf5* mutants (**M, N** respectively, white arrows, see also Supplemental movies 1 and 2).

To confirm that a single midline blood vessel is formed in *myod/myf5* mutants, we injected *mcherry-caax*.*p2a*.*nls-kikume* mRNA into wild type and mutant embryos to label the cell membranes and nuclei. Confocal microscopy shows that the wild-type embryos have two distinct blood vessels representing an artery and vein (Figure 1K). The *myod/myf5* mutants, however, showed one large vessel beneath the notochord (Figure 1L). This phenotype is not due to angioblast specification or migration defects, as specification and migration appear normal in *myod/myf5* morphants (Supplemental Figure 1B). Analysis of circulation at 25 hpf shows that circulation is completely lost in *myod/myf5* mutants compared to wild type (Figure 1M-N. Supplemental Movies 1 and 2)

### MyoD and Myf5 act upstream of Vegf and downstream of Hedgehog to induce the artery

A single midline vessel with a venous identity is consistent with disruptions in the Hedgehog-VEGF-Notch pathway that is required for arterial formation. Previous studies showed that loss of *vegfaa* causes a failure of dorsal aorta formation (Casie Chetty et al., 2017; Coultas et al., 2010; Lawson et al., 2002). To determine whether *vegfaa* expression in disrupted in *myod/myf5* double mutants, we performed an in-situ hybridization using an anti-sense probe against *vegfaa* and found that *vegfaa* expression was absent in *myod/myf5* double mutants (Figure 2A and 2C). Notch signaling has been shown to be downstream of VEGF signaling and required for arterial formation (Lawson et al., 2002, 2001) To confirm if Notch signaling was lost in the endothelium, we performed in-situ hybridization of Notch ligand *dll4*, an arterial specific Notch receptor. This showed a broad loss of expression in the endothelium of *myod/myf5* double mutants (Figure 2B and Figure 2D).

**Figure 2.**
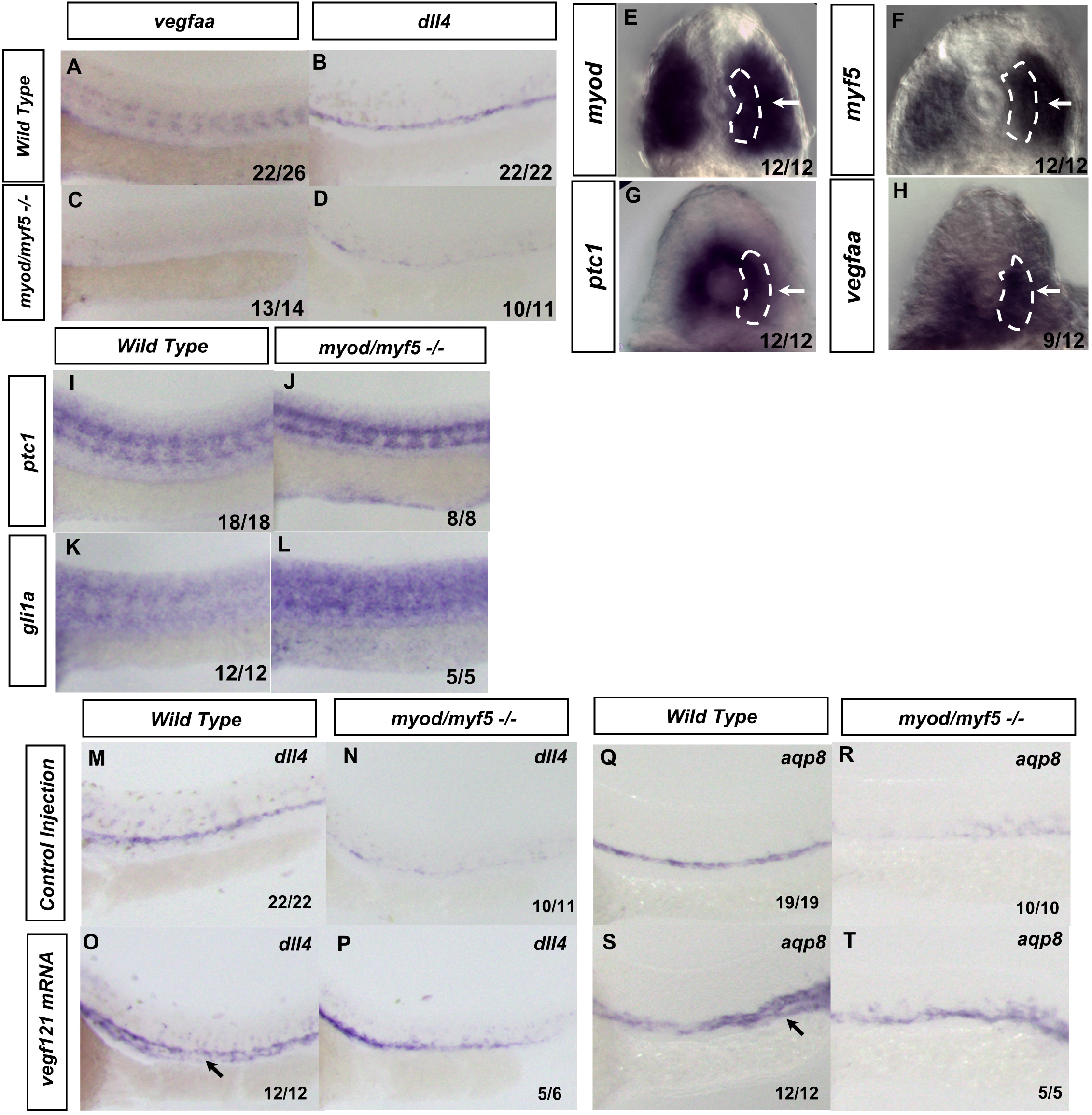
*myod/myf5* double mutants causes disruptions in the VEGF-Notch signaling pathway downstream of Hedgehog signaling. Whole-mount in-situ hybridizations of 25 hpf wild-type siblings (**A, B**) and *myod/myf5* -/- embryos (**C, D**). Wild-type embryos show clear expression of *vegfaa* (**A**) and *dll4* (**B**). In *myoD/myf5* mutants, both *vegfaa* and *dll4* are lost (**C, D**, respectively). DIC images of 19 hpf embryos, sectioned at the 5^th^ somite and stained for *myod* (**E**), *ptc1* (**F**), and *vegfaa* (**G**), or sectioned at the 15^th^ somite and stained for *myf5* (**H**). White arrows indicate somitic expression of indicated genes at overlapping locations. Whole-mount in-situ hybridization of 25 hpf embryos of hedgehog genes *ptc1* (**I, J**) or *gli1* (**K, L**). WT embryos (**I, K**) and *myod/myf5* mutants (**J, L**) show no change in *ptc1*. Interestingly, *gli1* appears to increase in the *myod/myf5* mutants (**L**). Whole-mount in-situ hybridization of 25 hpf embryos of Notch ligand *dll4* (**M-P**) and *aqp8* (**Q-T**). WT embryos (**M**,**O**,**Q**,**S**) and *myod/myf5* mutants (**N**,**P**,**R**,**T**) were injected with either *vegf121* mRNA (**O**,**P**,**S**,**T**) or vehicle (**M**,**N**,**Q**,**R**). Injections of *vegf121* into WT embryos induced an expansion of Notch *dll4* ligand and *aqp8a* arterial marker (**O**,**P**,**S**,**T**) compared to control injections (**M**,**N**,**Q**,**R**). In *myod/myf5* mutant embryos, injection of *vegf121* mRNA restored lost Notch and arterial markers. (**P**,**T**)

Previous work showed that Hedgehog signaling acts upstream of *vegfaa* to induce arterial formation (Coultas et al., 2010; Lawson et al., 2002; Wilkinson et al., 2012). We investigated if *myod* and *myf5* had overlapping expression domains within the somitic *hedgehog* receiving and *vegfaa* producing regions in the somite. We performed in-situ hybridization on *ptc1*, which encodes for a Hedgehog receptor and is a direct transcriptional target of Hedgehog (Concordet et al., 1996; Koudijs et al., 2008), along with *vegfaa, myod* and *myf5* on 19 hpf embryos (Figure 2J-2M). We then sectioned the embryos at the 5^th^ somite, except for the *myf5* which was sectioned at the 15^th^ somite as its expression domain is more posterior (Coutelle et al., 2001). The *myod* and *myf5* expression were localized in a pan somitic manner that included a medial somitic domain (Figure 2J, 2K, white arrows). *ptc1* and *vegfaa* were expressed in the medial somitic region proximal to the notochord (Figure 2L, 2M, white arrow and white dots), as well as the hypochord region, which lies ventral to the notochord. The overlapping expression domains of *ptc1* and *myod/myf5* in the medial somitic mesoderm indicate a convergence of Hedgehog and myogenic transcription factors that correspond to the *vegfaa* somitic expression domain.

To determine if *myod* and *myf5* loss of function affected hedgehog signaling components, we used probes for Hedgehog response genes *ptc1* and *gli1a* (Karlstrom, 2003; Karlstrom et al., 1999; Koudijs et al., 2008). In *myod* and *myf5* double mutants, *ptc1 and fli1a* are expressed normally in the somites (Figure 2O-R). Given that the somites maintain *gli1a* expression and that *ptc1* is a direct target of Hedgehog signaling, the results demonstrate that the Hedgehog pathway is still intact in *myod/myf5* double mutants (Chen and Struhl, 1996; Lewis et al., 1999).

To determine if *vegfaa* functions genetically downstream of *myod* and *myf5* during Notch signaling induced arterial differentiation, we attempted to rescue dorsal aorta formation in *myod/myf5* mutants by supplying exogenous *vegf* mRNA. We injected mRNA of the secreted form of *vegf, vegf121* (Liang et al., 2001) into *myod/myf5* double mutants and wild-type embryos to confirm if arterial fate would be restored (Figure 2S-2Z). After control vehicle injection, *myod*/*myf5* mutants contain little observable *dll4* or *aqp8a* in the trunk of the embryo (Figure 2T, 2X). However, after *vegf121* injection, wild-type embryos exhibit an expansion of *aqp8a* and *dll4* into ectopic regions of the vasculature (Figure 2U, 2Y, black arrows), and *dll4* and *cldn5b* were restored in *myod/myf5* mutants. (Figure 2V,2Z) This indicates that loss of the arterial fate in the *myod/myf5* mutants is the result of loss of *vegfaa* expression and subsequent absence of Notch signaling.

### Myod and Myf5 work in tandem with Hedgehog signaling to induce *vegfaa*

The Hedgehog signaling requirements for *vegfaa* expression have previously been determined through Hedgehog pathway mutants or by early stage treatment with Hedgehog inhibitors (Lawson et al., 2002), however it is unknown if Hedgehog signaling is continuously required for *vegfaa* expression after *myod* and *myf5* are expressed. To determine if this is the case, we administered the Hedgehog inhibitor cyclopamine at different stages of development. We utilized probes against *ptc1* to measure Hedgehog signaling output along with a probe against *vegfaa*. Embryos treated with control ethanol vehicle from 10-somite stage to the 22-somite stage showed robust *ptc1* and *vegfaa* expression (Figure 3A). However, cyclopamine treatment from the 20ss to the 22ss results in a reduced but not complete loss of both *ptc1* and *vegfaa* compared to wild type. (Figure 3B) Inhibition of Hedgehog signaling beginning at the 15ss or 10ss results in complete loss of somitic expression of *vegfaa* as well as somitic expression of *ptc1* at the 22ss (Figure 3C and 3D). *myod* and *myf5* expression were the same across all treatment conditions, indicating Hedgehog signaling is not reducing bHLH activity during treatment (Figure 3A-3D). Given that *vegfaa* expression corresponds closely to *ptc1* expression, it appears that Hedgehog signaling is continuously required for expression of *vegfaa* along with *myod* and *myf5*.

**Figure 3.**
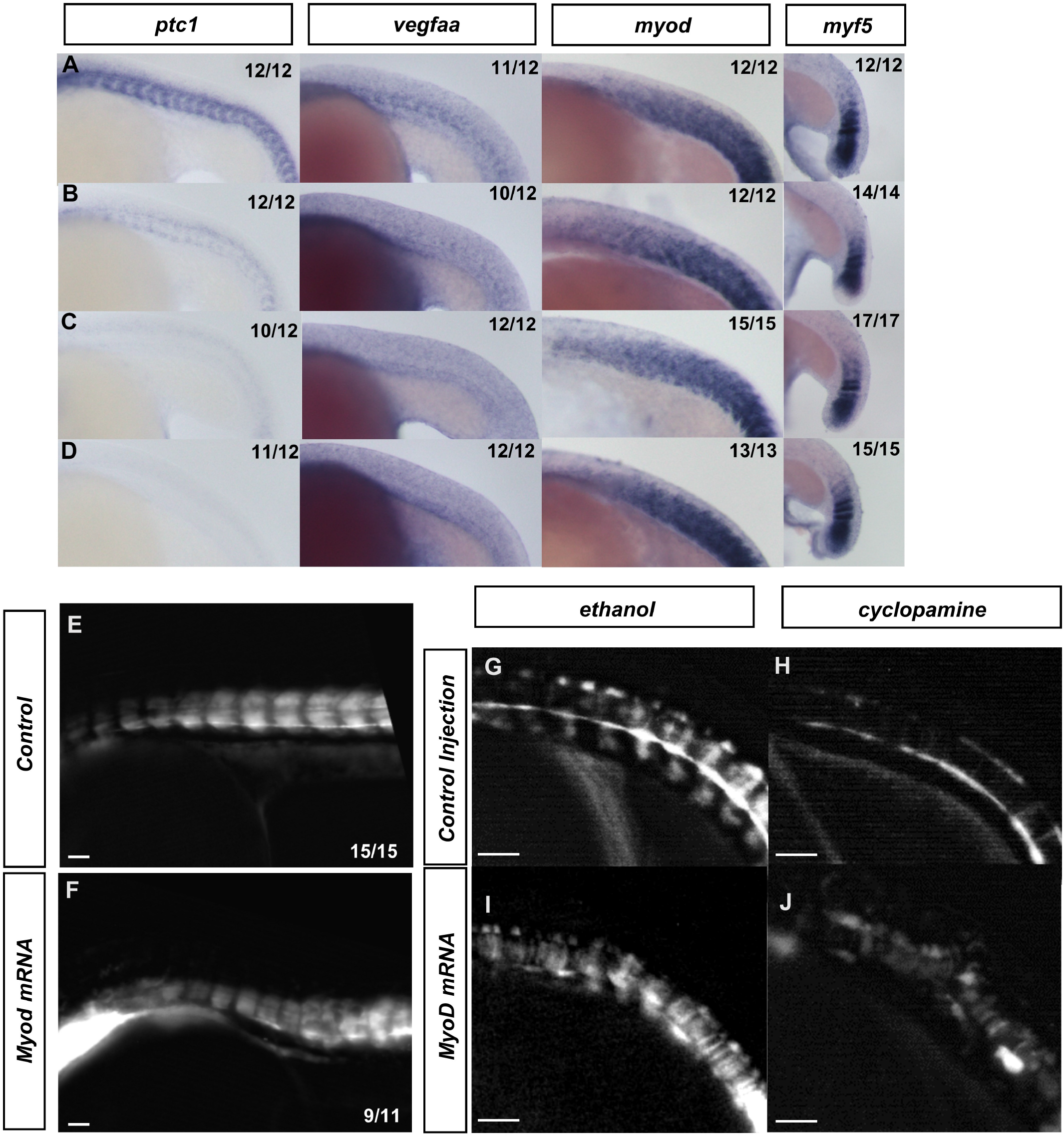
MyoD and Hedgehog signaling work cooperatively to induce somitic *vegfaa. bHLH* transcription factors are not sufficient to expand *vegfaa*. (**A-D**) In-situ hybridization of wild-type embryos treated with (**A**) Ethanol vehicle or (**B**) Cyclopamine from 20-somite stage to 22-somite stage, (**C**) from 15-somite stage to 22-somite stage, or (**D**) from 10-somite stage to 22-somite stage. In-situ probes were used against *ptc1, vegfaa, myod, and myf5*. 25 hpf *tg(vegfaa;GFP)* embryos were injected with vehicle (**E**) or *myoD* mRNA (**F**). 18-somite stage *tg(vegfaa;GFP)* embryos were treated with ethanol (**G**,**I**) or cyclopamine (**H**,**J**) from tailbud stage. These embryos were either injected with vehicle (**G**,**H**) or *myod* mRNA (**I**,**J**). White arrows indicate somitic expression of *tg(vegfaa:gfp)*.

To test if ectopic *myod* mRNA is sufficient to expand the *vegfaa* expression domain, we injected *tg(vegfaa:gfp)* embryos with *myod* mRNA to observe if there is an expansion of *vegfaa* expression. Vehicle injected embryos showed stereotypical *vegfaa* reporter expression (Figure 3E). Embryos injected with *myod* mRNA did not show expanded expression compared to vehicle injection (Figure 3F).). Interestingly, *myod* mRNA did not increase somitic *tg(vegfaa:gfp)* expression despite expansion of muscle reporter *tg(actc1b:gfp)* in injected embryos (Supplemental Figure 2).

While previous studies have shown the Hedgehog signaling is required for somitic *vegfaa* expression, it is also required for muscle differentiation and expression patterns of *myod* and *myf5* in the adaxial region (Lawson et al., 2002; Swift and Weinstein, 2009). To understand if Hedgehog induced *vegfaa* expression could be functioning through myogenic transcription factors, we injected *myod* mRNA into *tg(vegfaa:gfp)* embryos treated with Hedgehog inhibitor cyclopamine, or ethanol vehicle (Figure 3G-3J). Comparison of cyclopamine treated embryos injected with either *myod* or control showed no rescue of somitic *tg(vegfaa:gfp)* (Figure 3H,3J This indicates *myod* is not sufficient to drive *vegfaa* in the absence of Hedgehog.

To determine if Myod and Myf5 activity are continuously required for *vegfaa* expression, we utilized a heat shock inducible construct of the HLH protein ID3 driven by the hsp70 promoter (Row et al., 2018). This transgenic zebrafish known as *tg*(*hsp70:id3-p2a-nls-kikume)* (hereafter referred to as *HS:Id3)*, inhibit that activity of *myod* and *myf5* transcription factors by competitively suppressing their E-protein binding partners (Row et al., 2018). Hemizygous *HS:Id3* adults were crossed to either *tg(vegfaa:gfp)* or *tg(actc1b:gfp)* and were heat shocked at the 8-somite stage. Embryos were grown to 30 hpf and *kikume* was photoconverted from green to red with 405 nm light prior to imaging. The *tg(actc1b:gfp)* reporter labels differentiated skeletal muscle (Figure 4A,4B). White arrows indicate areas of GFP depletion as a result of ID3 overexpression. Loss of GFP from *tg(actc1b:gfp)* is restricted to posterior regions of the embryo, with GFP disappearing around the 12^th^ somite and returning in the tail of the embryo, presumably after the transient expression of Id3 is depleted (Figure 4B). On the other hand, loss of GFP from *tg(vegfaa:gfp)* includes anterior and posterior regions of the embryo, with GFP returning in the tail of the embryo (Figure 4D). To quantify which regions specifically retained *tg(vegfaa:gfp)* reporter expression, we measured GFP expression in 3 sectors of the embryo with roughly equivalent somite numbers: the anterior 10 somites, the following 10 somites, and the most posterior somites (Figure 4E). Wild type somites express *tg(vegfaa:gfp)* throughout all sectors, with mild variability in the posterior somites as they vary in number. Following *HS:Id3* activation, *tg(vegfaa:gfp)* expression in the first two sectors is diminished, while the third retains activity (Figure 4E). This domain of GFP activity can be altered depending on the stage of heat shock, as later induction at 15-somite stage specifically inhibits expression in the posterior sector (Supplemental Figure 3B and 3D, respectively). This indicates that there is a window of bHLH activity that is required in somites to maintain expression of *vegfaa*. When exogenous Id3 is depleted after the transient heat shock, *vegfaa* expression can return in newly formed somites.

**Figure 4.**
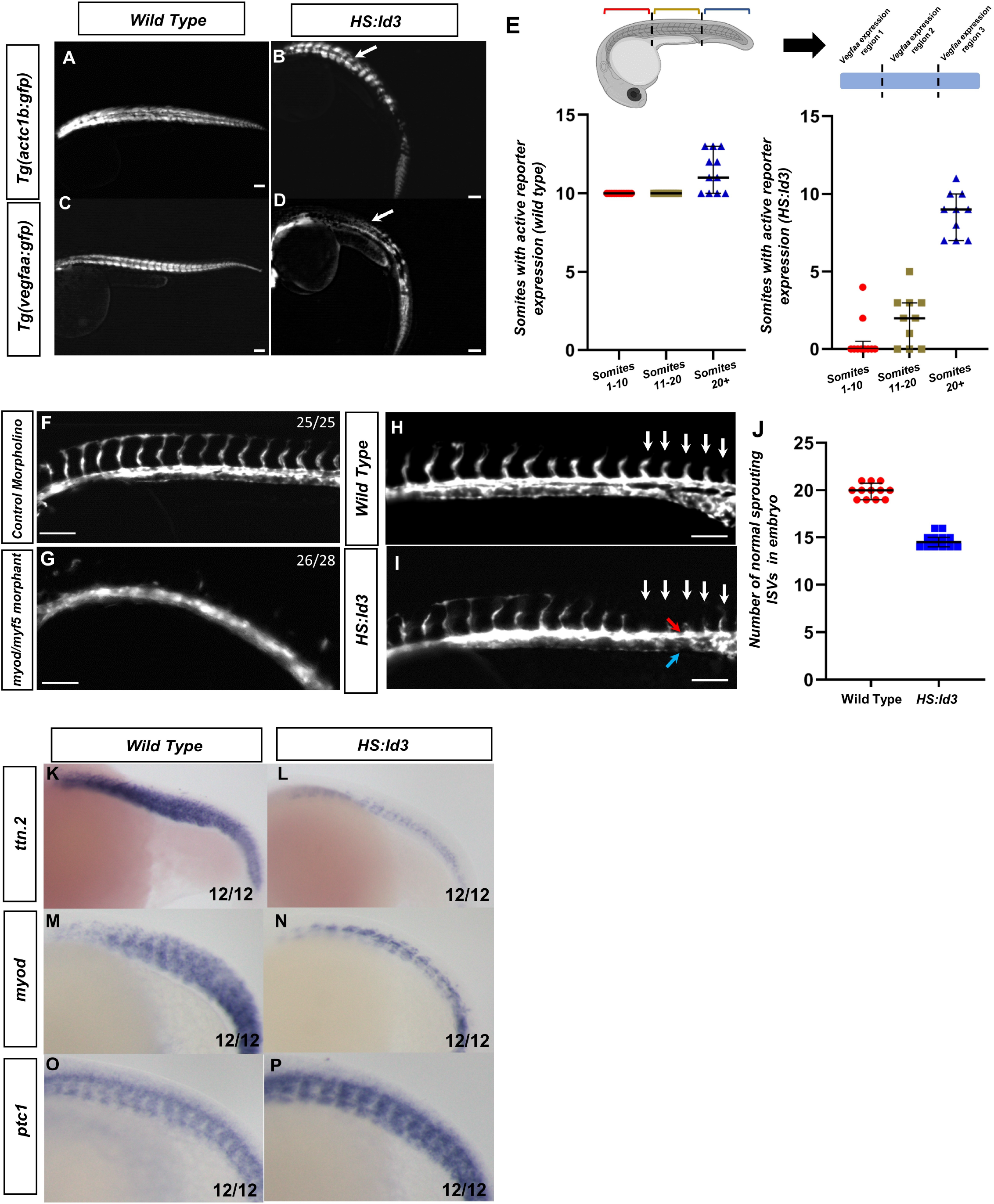
bHLH transcriptions factors are required after somitic mesoderm and muscle specification for *vegfaa* expression. Fluorescent images were taken of 30 hpf embryos with the following genotype: *tg(actc1b:gfp)* (**A**), *tg(actc1b:gfp)/HS:Id3* (**B**), *tg(vegfaa:gfp)* (**C**), or *tg(vegfaa:gfp)/HS:Id3* (**D**). All embryos were heat shocked at 38.5°C for 30 minutes at the 8-somite stage. Fluorescent images of *HS:Id3* show loss of anterior GFP signal in *tg(vegfaa:gfp)* (**D**), but not *tg(actc1b:gfp)* (B). (**E**) Quantification of the number of somites with active *tg(vegfaa:gfp)* expression in either control (N=11) or *HS:id3* (N=10) embryos. Regions were split into somites 1-10, somites 11-20, and expression after somite 20+. The P-values for each somite sector are *p>*.*0001, p>*.*0001*, and *p=*.*0013* for section 1, 2, and 3 respectively. (**F-G**) Florescent images of morphant *tg(kdrl:gfp)* embryos showing ISV formation in control (**F**) and in absence of *myod/myf5* (**G**). (**H-I**) Fluorescent images of vascular reporter showing sprouting ISVs in (**H**) *tg(kdrl:egfp or* (**I**) *tg(kdrl:egfp)/HS:Id3* (red arrow and blue arrow indicate dorsal aorta and cardinal vein, respectively). (**J**) Quantification of normal sprouting ISVs in wild type and *HS:Id3* embryos (N=12 for each condition, P<.0001). Whole-mount in-situ hybridization of *HS:Id3* (**L**,**N**,**P**) and wild type siblings (**K**,**M**,**O**) were done on 20-somite stage embryos with following probes: *ttn*.*2* (**K**,**L**) *myod* (**M**,**N**) and *ptc1*(**O**,**P**). In-situ hybridizations of *ttn*.*2, myod*, and *ptc1* show the anterior loss of *tg(vegfaa:gfp)* is not correlated with total loss of differentiated muscle or Hedgehog signaling. (**Q**) Schematic showing that *myod* and *myf5* are required to generate myogenic mesoderm, which work cooperatively with Hedgehog to induce Vegf secretion.

Embryos that lack *myod/myf5* fail to form intersegmental vessels (ISVs) when compared to wild type siblings, a phenotype consistent with VEGF signaling loss (Figure 4F and 4G, respectively. Supplemental Figure 4). To determine if *bHLH* transcriptions factors are required continuously for the development of the endothelium and ISVs, we utilized the *HS: Id3* line to deplete *bHLH* binding partners required for their function. The *HS:Id3* line was crossed to *tg(kdrl:eGFP)* to label to the blood vessels. The embryos were heat shocked at the 15-somite stage, to allow for the dorsal aorta to form properly as VEGF is dispensable following this stage (Casie Chetty et al., 2017), and grown to 24 hpf. Analysis of the embryos shows that ISVs sprout normally throughout the trunk of control embryos (Figure 4H). However, the ISVs for *HS:Id3* embryos show impaired ISV formation, with ISVs failing to sprout correctly after the 13th somite (Figure 4I). The difference between the number of control ISVs and *HS:Id3* ISVs showed a failure to form the correct number of sprouting ISVs, even though the dorsal aorta was able to form correctly and distinctly from the cardinal vein (Figure 4J, red arrow indicates dorsal aorta and blue arrow indicates the cardinal vein). Interestingly, *vegfaa* promoter activity seems to be immediately compromised throughout the embryo after heat shock, as indicated by *tg(vegfaa:gfp)* embryos losing *gfp* transcript in the *HS:Id3* background (Supplemental Figure 5). However, ISVs are not lost in the anterior region, indicating there is a temporal requirement of bHLH activity for *vegfaa* induced sprouting to drive ISV formation.

In order to confirm that *HS:Id3* activation did not affect a cell fate decision or Hedgehog signaling, in-situ hybridizations were performed on *HS:Id3* embryos and wild type siblings on embryos that were heat shocked at 8ss. Probes for *ttn*.*2, myod*, and *ptc1* were used to label differentiated muscle, *myod* expressing cells, and Hedgehog signaling activity, respectively (Figure 4K-4P). While differentiated muscle and *myod* expression was reduced in *HS:Id3* embryos, they was not completely lost in the anterior regions of the embryo (Figure 4L and 4N). However, Hedgehog signaling was largely unaffected in *HS:Id3* embryos, as indicated by broad *ptc1* expression (Figure 4P). This shows that loss of *tg(vegfaa:gfp)* signal was not due to a loss of Hedgehog signaling or a complete absence of muscle tissue..

In this study, we show that *myod* and *myf5*, along with Hedgehog signaling, act upstream of *vegfaa* to induce dorsal aorta formation. This is the first time this connection between MRFs and non-autonomous differentiation has been shown in an embryological context. Previous studies examined similar phenomena in cultured myoblasts or in adult skeletal muscle tissue, where differentiating myofibers can secrete Vegf. (Bryan et al., 2008; Chiristov et al., 2007; Renault et al., 2013; Verma et al., 2018) Our study shows a surprising role for myogenesis that places it upstream of midline endothelial patterning in the zebrafish embryo. The requirement of myogenesis for Vegf induction indicates that Hedgehog alone is not sufficient for *vegfaa* expression. Indeed, Hedgehog signaling is still robust in the mesoderm even in the absence of *myod* and *myf5*. However, it does not appear that *myod* and *myf5* expression colocalizes with *vegfaa* entirely, nor can exogenous *myod* rescue *vegfaa* expression when Hedgehog is lost.

We therefore propose that Hedgehog and MRFs act in tandem to induce *vegfaa*, with midline derived Hedgehog signaling cooperating with MyoD and Myf5 to induce *vegfaa* in the medial somite. Indeed, both the dorsal aorta and their ISVs develop adjacent to the medial somite where *vegfaa* is expressed (Jin et al., 2005). We also showed that bHLH transcription factors and Hedgehog signaling are continuously required, even after somite formation, for *vegfaa* expression maintenance. This implies that Hedgehog and Myod and Myf5 continue to cooperate to maintain *vegfaa*. Interestingly, previous studies found synergistic effects of Gli transcription factors with multiple transcriptional pathways, including bHLH TFs in particular (Elliott et al., 2020; Lee et al., 2010; Sabol et al., 2018). It is possible Gli, with MyoD and Myf5, work in physical proximity in the genome to generate tissue specific expression. Further research is required to determine if a functional interaction of the proteins themselves is required for *vegfaa*.

## MATERIALS AND METHODS

### In Situ Hybridization and Immunohistochemistry

Whole-mount in situ hybridization was performed as previously described (Griffin et al., 1995). Antisense RNA probes were made for *stab1l* (Rost and Sumanas, 2014), *aqp8a* (Sumanas et al., 2005), *etv2* (Sumanas et al., 2005), *fli1a* (Thompson et al., 1998) *gli1a* (Karlstrom, 2003), *ptc1* (Concordet et al., 1996) *ttn*.*2* (Maves et al., 2009). The *cldn5b, dll4*, and *dab2* DIG probes were generated by *taq*-generated PCR fragments integrated into a PCRII vector using the Topo-TA Cloning kit (Thermo Fisher). *vegfaa* probe was generated from *vegf165* pCS2+ vector (Sumanas and Lin, 2006).

### Microinjections

mRNA was prepared from full length *shh* and *vegf121* clones in the pCS2+ vector. They were linearized with NotI enzyme, and transcribed using an SP6 mMessage Machine kit (Ambion). *myod* mRNA was prepared by cloning a gBlock synthesized with the *myod-3xflag* open reading frame into a PCS2+ vector (Integrated DNA Technologies). The plasmid was linearized using KpnI enzyme and transcribed using SP6 mMessage Machine kit. *mCherry-Caax-nls-kikume* mRNA was generated from a NotI enzyme linearized *HS:mcherry-caax-nls-kikume* plasmid described previously (Goto et al., 2017). One-cell stage zebrafish embryos were injected with 50 picograms of *vegf121* mRNA, 100 picograms of *mcherry-caax-nls-kikume* mRNA, 35 picograms *shh* mRNA, or 30 picograms of *myod* mRNA. 2.5 ng each of *myod* and *myf5* morpholino, or 5.0 ng of control morpholino, were injected into one cell stage embryos as described previously (Row et al., 2018).

### Microscopy and Imaging

DIC time-lapse images and fluorescent images were performed using a Leica DMI6000B inverted microscope with a 10× objective. In-situ hybridization sections were imaged using DIC on the Leica DMI6000B inverted microscope with a 40× objective. Bright-field images of whole mount in-situ hybridizations were obtained using a M165FC microscope (Leica) equipped with an Infinity 3 camera (Lumenera). Fluorescent images for sections (Figure 2) were imaged on a custom assembled spinning disk confocal microscope consisting of an automated Zeiss frame, a Yokogawa CSU-10 spinning disc, a Ludl stage controlled by a Ludl MAC6000 and an ASI filter turret mated to a Photometrics Prime 95B camera. This microscope was controlled with Metamorph microscope control software (V7.10.2.240 Molecular Devices), and laser power levels were set in Vortran’s Stradus VersaLase 8 software. Images were processed in ImageJ.

### Zebrafish drug treatments

Hedgehog was inhibited by cyclopamine. Cyclopamine treatments were performed from stock solutions of 20 mM cyclopamine (Sigma) in ethanol diluted into 20 µM working solutions in embryo media.

### Zebrafish Lines

The *myod*^*fh261*^ and *myf5*^*hu2022*^ mutant strains were maintained on the AB background and were previously described (Hinits et al., 2011, 2009). Compound heterozygous mutants for *myod*^*fh261*^ and *myf5*^*hu2022*^ alleles were generated by in-cross and grown to adulthood. Genotyping was done as described previously (Row et al., 2018). *tg(kdrl:eGFP)* ^*s843*^, *tg(actc1b:gfp)* ^*zf13tg*^, *tg(hsp70l:id3-2A-NLS-KikGR)* ^*sbu105*^, and *tg(hsp70l:CAAX-mCherry-2A-NLS-KikGR)* ^*sbu104*^ lines were maintained as done previously (Goto et al., 2017; Higashijima et al., 1997; Jin et al., 2005; Row et al., 2018). *tgBAC(vegfaa:EGFP)* ^*pd260Tg*^ was maintained as hemizygotes (Karra et al., 2018). For Hedgehog manipulated embryos (Figure 3 and Figure 4), manipulated embryos are a TLB background used previously (Goto et al., 2017).

## Acknowledgements

We thank Stephanie Flanagan for excellent fish care. We thank Miguel Torres-Vasquez and Kenneth Poss for providing the *tgBAC(vegfaa:EGFP)* ^*pd260Tg*^ transgenic line, and Lisa Maves for providing the *myod*^*fh261*^ and *myf5*^*hu2022*^ mutant lines. The *vegf165* and *stab1l* plasmids were generously gifted to us by Dr. Saulius Sumanas, and the *ptc1* and *gli1a* probes were generously gifted to us by Dr. Howard Sirotkin. We also thank Dada Pisconti for critical reading of the manuscript.

