## Supplemental figures and legends for "Myogenic regulatory factors MyoD and Myf5 are required for dorsal aorta formation and angiogenic sprouting"

Stony Brook University

Stony Brook, NY 11794-5215

**Supplemental Figure 1. Myod and Myf5 are not required for angioblast migration**

Time-lapse fluorescent images were taken of 10-somite stage tg(kdrl:eGFP) embryos with the following morpholinos: Control (A) or *myod* and *myf5* (B). Images were taken every sixty minutes for 180 minutes. White arrows show angioblasts migrating to the midline.

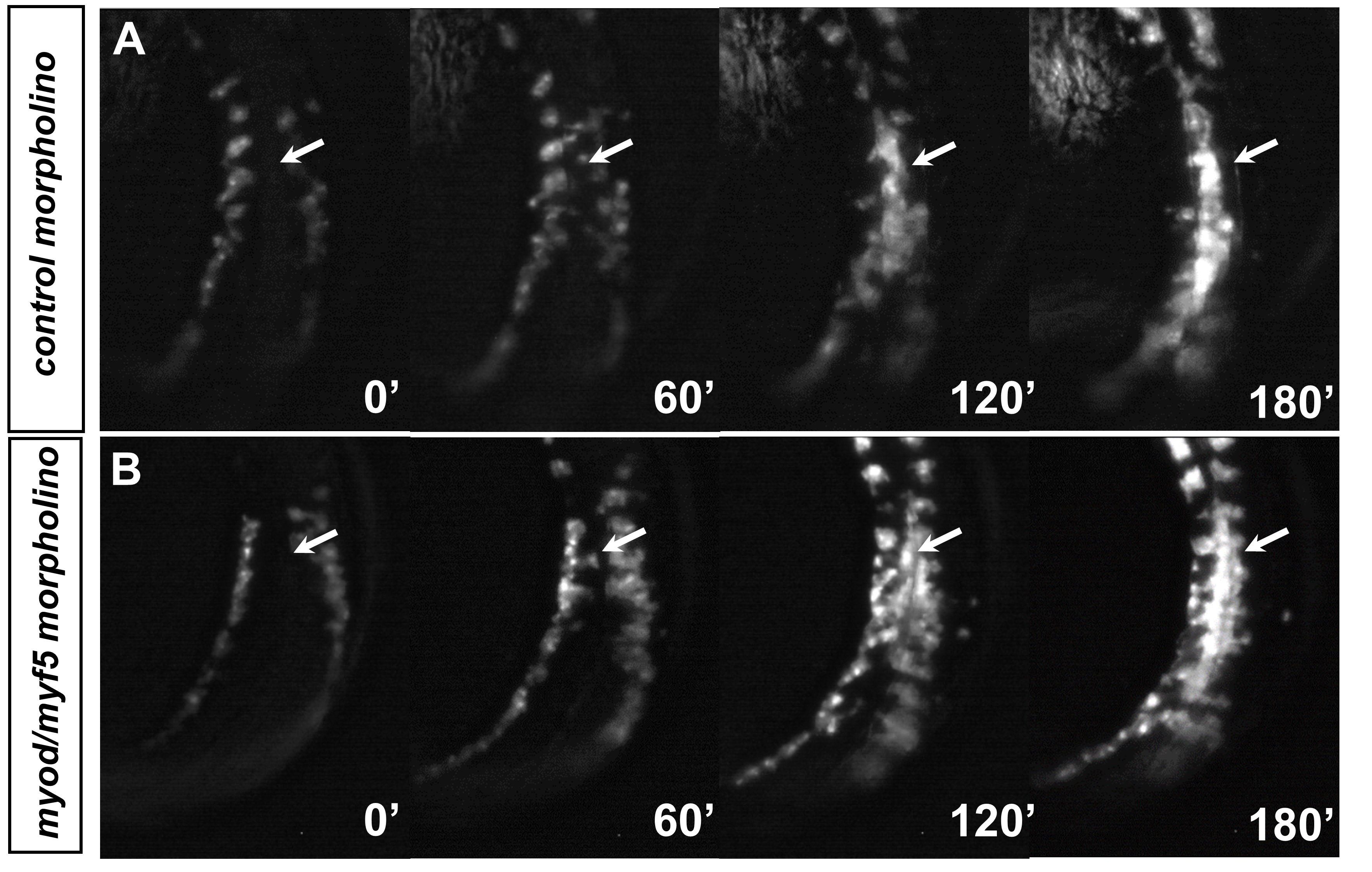

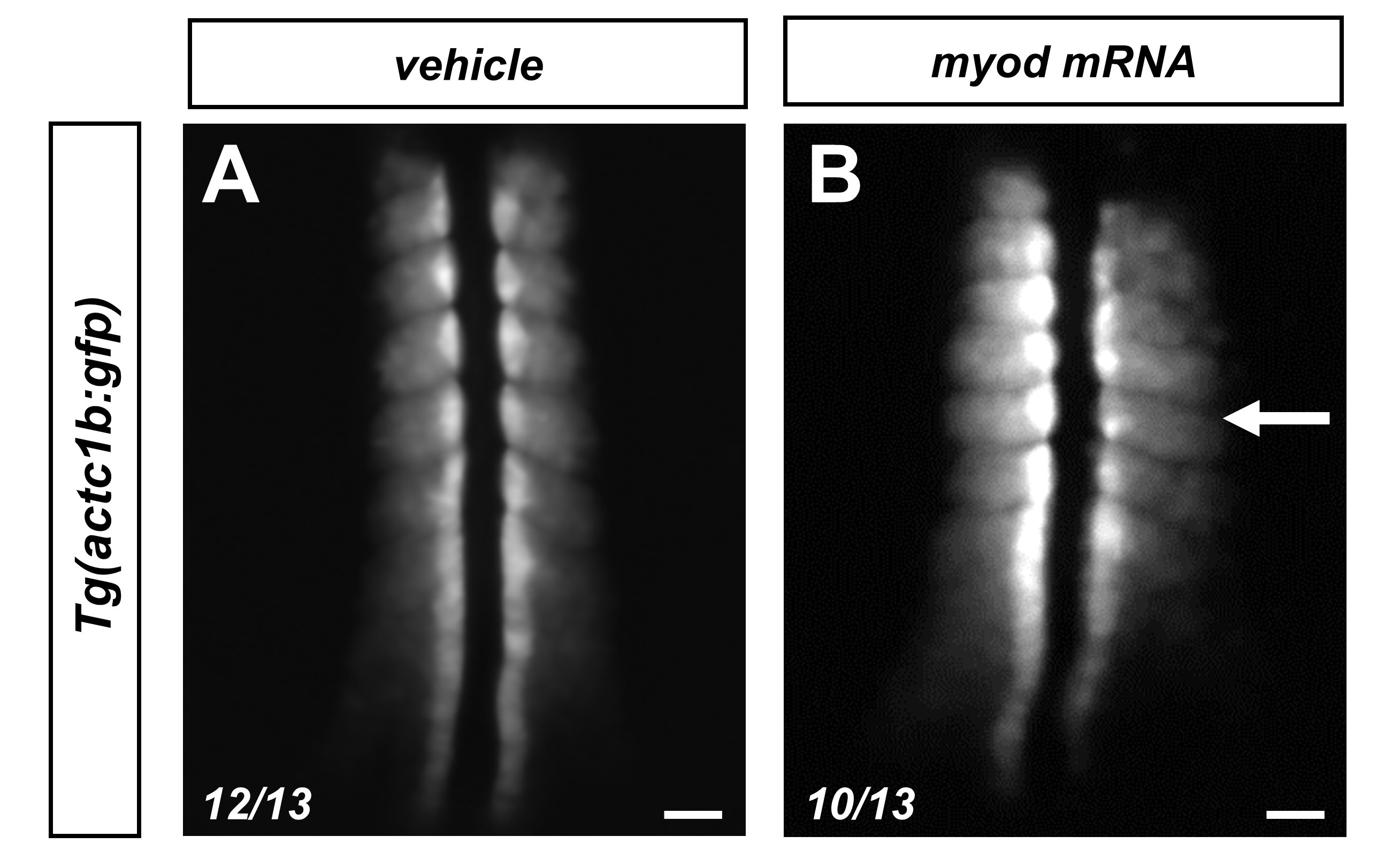

**Supplemental Figure 3. Late stage bHLH inhibition shifts *vegfaa* expression loss posteriorly.**

Fluorescent images were taken of 30 hpf embryos with the following genotype: *tg(actc1b:gfp)* (**A**), *tg(actc1b:gfp)/HS:Id3* (**B**),  *tg(vegfaa:gfp)* (**C**), or *tg(vegfaa:gfp)/HS:Id3* (**D**). Individual somites were quantified for expression of indicated transgenic reporters (**E**). All embryos were heat shocked at 38.5°C for 30 minutes at the 15-somite stage. White arrows show lost reporter activity in the posterior regions of the embryo.

**Supplemental Figure 2. *myod* injected embryos show expanded muscle**

*tg(actc1b:gfp)* embryos imaged at the 6-somite stage were injected with vehicle (A) or *myod* mRNA (B).

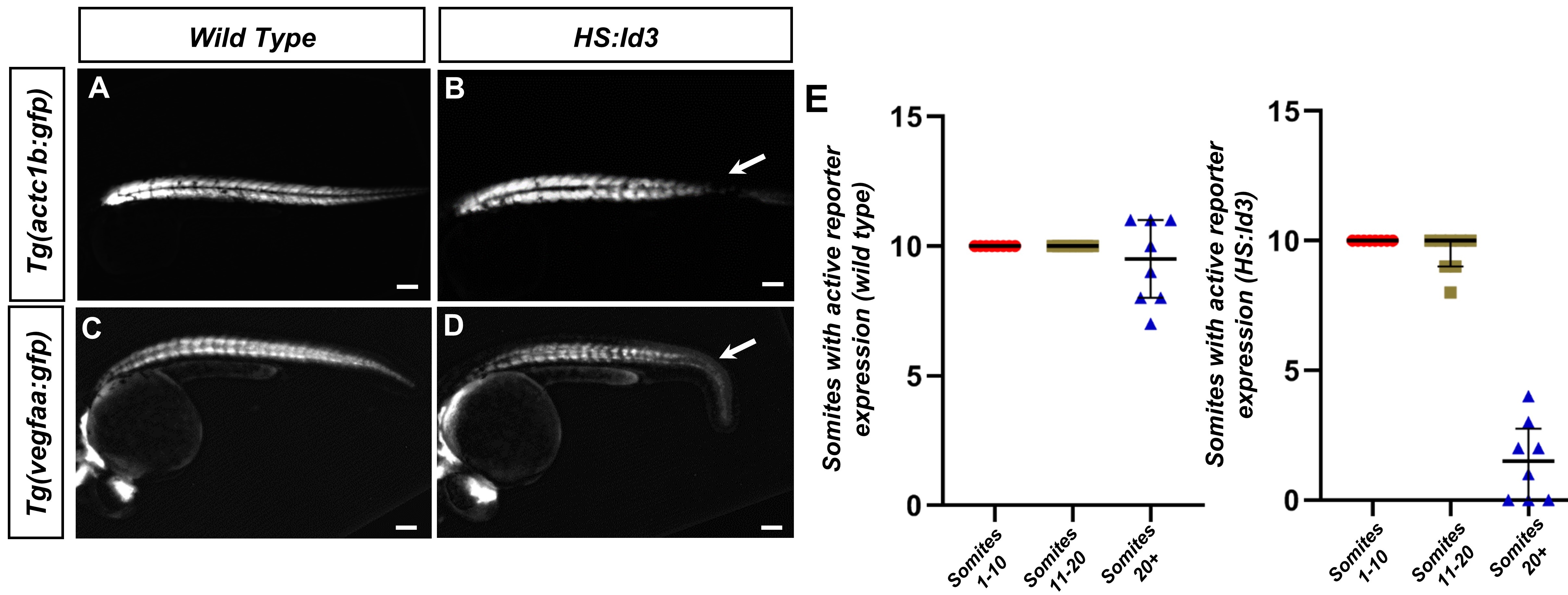

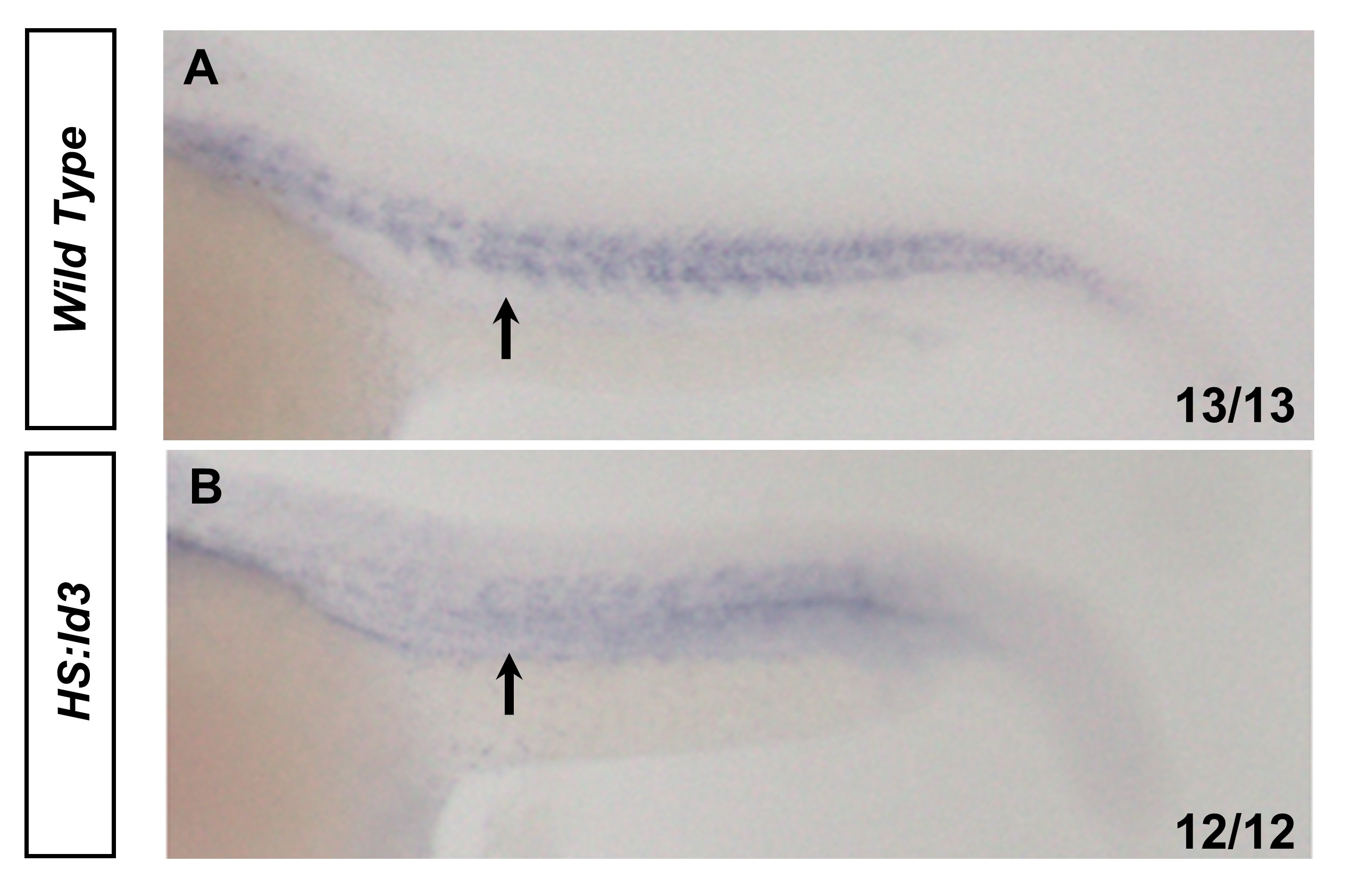

**Supplemental Figure 4. *myod/myf5* mutants show no clear ISVs**

In-situ hybridizations for *fli1a* show ISVs in vasculature of (**A**) *wild type* but not (**B**)  *myod/myf5 -/-.*

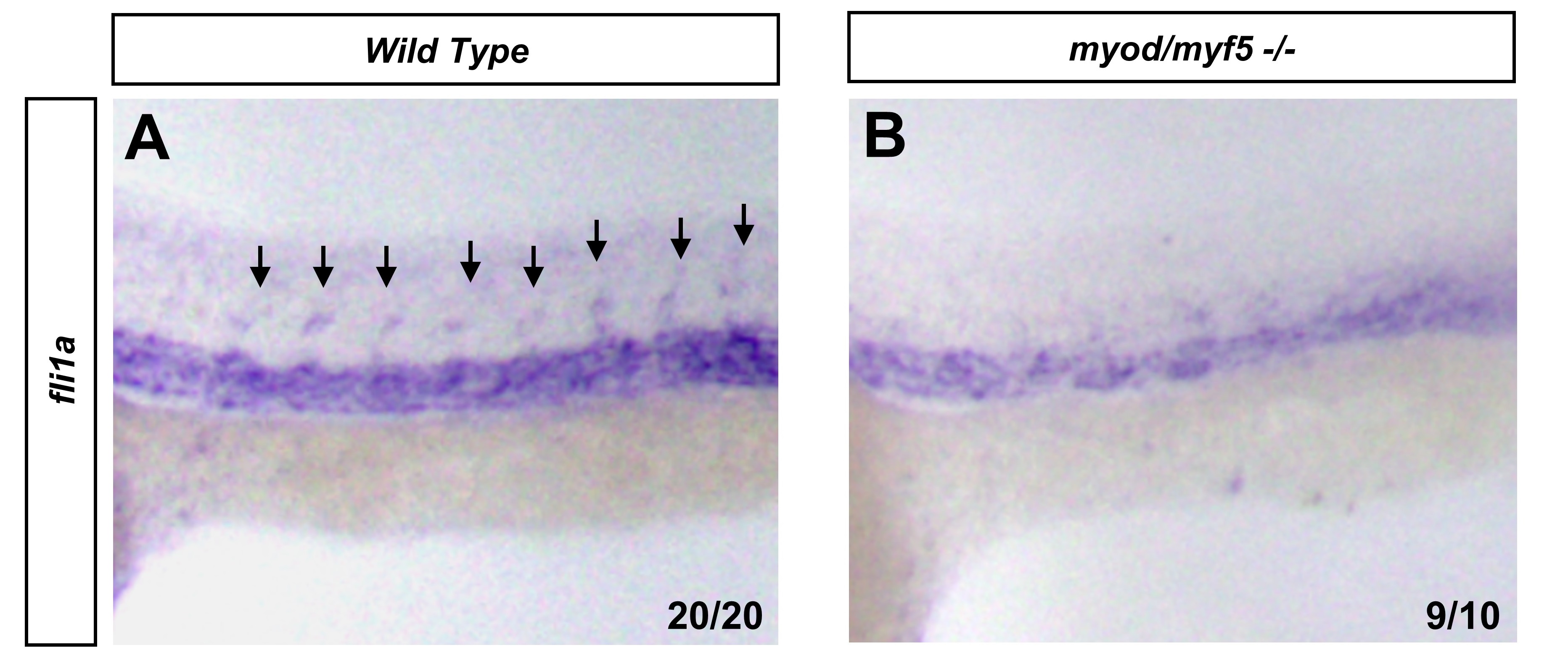

**Supplemental Figure 5. Loss of bHLH activity results in immediate loss of *vegfaa* expression**

Whole-mount in-situ hybridization *tg(vegfaa:gfp)* embryos in either wild-type (**A**) or *HS:Id3* (**B**) background with *gfp* probe at 24 hpf. Embryos were heat-shocked at the 15-somite stage. Black arrows show somitic *gfp* expression.

**Supplemental Movie 1. Normal circulation of wild-type embryo.**

A DIC movie showing blood cells circulating through an embryo over a 25 second period.

**Supplemental Movie 2. Absent circulation of *myod/myf5* double mutant embryo.**

A DIC movie showing double mutant embryos through an embryo over a 25 second period. Note the lack of blood flow.
